# Mitochondrial maintenance is involved in the exceptional longevity of reproductive queens of the eusocial ant *Lasius niger*

**DOI:** 10.1101/2024.06.27.600950

**Authors:** Maïly Kervella, Fabrice Bertile, Alexandra Granger-Farbos, Benoît Pinson, Alain Schmitt, Martin Quque, Frédéric Bouillaud, François Criscuolo

## Abstract

Most social insects are characterized by a wide disparity in life-history traits between individuals of the same species. Sterile workers live for months or years while queens may live for decades. Theories of ageing emphasise the importance of metabolism and oxidative stress in explaining longevity, with mitochondrial bioenergetics standing at the crossroads of energy and reactive oxygen species production. Studying mitochondrial functioning therefore takes on its full relevance in determining the nature of the mechanisms that explain the contrasting longevities between insect social castes. We addressed this question in an eusocial species, the black garden ant *Lasius niger*. We found that caste differences in mitochondrial bioenergetics and oxidative balance only partially match with predictions of the oxidative stress theory of ageing. Long-lived queens were characterized by a lower metabolic rate, lower mitochondrial density yet not necessarily lower levels of mitochondrial oxidative damages. Despite this, queens did not show reduced ATP content; rather, they even possessed a higher energy load in their mitochondria. Converging clues suggested better mitochondrial maintenance in queen ants, with enhanced dynamics of mitochondrial fission and fusion and a more marked expression of mitochondrial enzymes of the Krebs cycle. Overall, our data paves the way for studying deeper into how the rate of ATP production *per* mitochondria is related to the investment in mitochondrial and somatic cellular maintenance, and whether it has specifically been selected as a key mechanism in defining the still unexplained paradoxical longevity of the queen reproductive caste.

## Introduction

In the fields of gerontology and anti-ageing medicine, a distinction is made between chronological age, which is the amount of time elapsed since birth, and biological/or physiological age, estimated using the progressive and highly variable rate of accumulation of damages in biomolecules and somatic cells. They lead to compromised biological functions proposed as hallmarks of ageing [1]. The list of the original 9 hallmarks (among which genomic instability, telomere attrition, mitochondrial dysfunction, cellular senescence…) recently completed by additional ones [2](e.g. compromised autophagy, inflammation) underline the involvement of mitochondria as the main site of both generation and integration of oxidative stress in the ageing process [3–5].

Mitochondria stand at the crossroads of bioenergetics and oxidative stress theory of ageing since: 1) this organelle is often referred to as the “powerhouse” of the cell because it is primarily responsible for producing adenosine triphosphate (ATP), the cell’s main source of energy, 2) mitochondria are also referred to as the main producers of inevitable by-products of aerobic metabolism, namely reactive oxygen species (ROS) [6–8]. In that respect, some old-established theories of ageing emphasise the importance of energy metabolism and oxidative stress in explaining longevity and senescence [9,10,6]. Ageing is typified by a progressive decline in mitochondrial activity and in mitochondrial stress response, leading to multiple effects such as decreased energy production (ATP), enhanced ROS production, accumulation of deleterious mitochondrial DNA mutations and abnormal accumulation of metabolites of the tricarboxylic acid cycle [11,12]. Some of these hallmarks of mitochondrial ageing have been key instruments in triggering the study of mitochondria in the field of the evolutionary biology of ageing [13,14]. Evolutionary biologists are not interested in solving the ageing issue, but rather in explaining the origins of the variability in ageing rates among and within species. In that context, an ever-growing numbers of studies have paved the way for mitochondrial explanations of, from the interspecific to the interindividual levels of interest, the paradoxical longevity of hypermetabolic birds [15,16], the relationship between mitochondrial respiration and the whole organism energy expenditure [17,18], and as a correlate the link with individual fitness through the regulation of energy production and investment in competing life-history traits such as growth, reproduction and somatic maintenance (i.e., longevity) [19,20]. To date, the large majority of these studies implemented the mitochondria in their explanatory schema by focusing on the multiple links among mitochondrial respiration (O_2_ consumption) and ATP/ROS production, using mitochondrial efficiency (i.e., the rate of ATP per O_2_ consumed or ROS produced) as the key variable [21–23]. In a nutshell, the oxidative phosphorylation (OXPHOS) merges (i) the creation of a protonmotive force between the mitochondrial matrix and the internal mitochondrial membrane space using the oxidation of food-derived energy into electron transport by the electron transport chain (i.e., OX), and (ii) the phosphorylation of ADP into ATP by the ATP synthase that feeds on the protonmotive force (i.e., PHOS). The coupling between these OX and PHOS components remains variable and under the modulation by internal (e.g., internal membrane protons’ leaking, *i.e.* not returning in the mitochondrial matrix through the ATP synthase, [24]) and external factors (such as diet, [25]). Thus, measuring mitochondrial efficiency has proved to be of primary importance in enlightening us about the mechanisms underlying life-history trade-offs in multiple species [26,27]. However, the implication of mitochondrial bioenergetics in life-history trade-off and ageing cannot be apprehended solely through the change in mitochondrial efficiency (coupling) over age since (i) mitochondria respond not only to the ATP demand but also in relation to substrate availability (ADP) [28], (ii) the ratio ATP/ADP is likely to be a regulator of the rate of ATP hydrolysis [29], and as such of the amount of energy invested in somatic maintenance, (iii) mitochondria may affect cell ageing via particular maintenance processes like the fission-fusion mechanism [30]. For instance, mitophagy, i.e., the removal of non-functional mitochondria and thereby the maintenance of an efficient aerobic metabolism (ATP produced at low ROS-related cost), is reduced throughout the ageing process in animals [2].

Social insects are a unique research animal model, notably because genetically similar individuals can express distinct phenotypes associated with different ageing rates [31]. In comparison to non-social species, they are notably characterized by immutable sterile castes (extreme altruistic behaviour), leading to a division of labour that defines eusocial species [32]. In eusocial Hymenoptera (ants, bees and wasps), at least two distinct female castes coexist with specific functions: a single or a few queens, dedicated to reproduction, and workers, which can be either entirely (for the eusocial species) or partially sterile and carry out non-reproductive tasks. In this case, female breeders can live longer than the non-breeding individuals, with ant queens living by one or even two decades longer than workers, whereas males die soon after the mating flight [33] (see also [34,35] for termites). Thus, the usual life history trade-off between fecundity and longevity does not seem to apply to social insects [13,32] and a fundamental challenge is to understand how a single genome can produce alternative phenotypes, and particularly alternative ageing phenotypes in monogyne species (i.e., all individuals originating from a single breeder). *Lasius niger* is a monogyne ant species, where the queen and the sterile workers coexist [33,36]. Within the latter, there is a division of labor based on temporal polyethism, with the specific behaviour of workers depending on their chronological age: older individuals are in charge of the riskiest task involving leaving the nest in search for food, while younger workers concentrate on brooding and caring for the queen. The lifespan of *Lasius niger* queens is about 10 times that of workers [37,38]. Thus, despite being genetically very close, the rate of biological ageing is extremely dissimilar within a colony, depending on individual’s social and behavioural status.

Mechanistic data on ageing in social insects actually indicate caste specificities. In honeybees, functional decline of brain or motor functions reflects lifespan disparities between workers of different castes, with foragers presenting cues of reduced learning performances for a given chronological age [39,40]. Lipofuscin is a pigment that accumulates in multiple somatic cells with age and has been related to age-related diseases including brain disorders [41,42]. In social insects, lipofuscin accumulates at a lower rate in tissues of long-lived worker honeybees than in short-lived individuals of the same age [39]. Therefore, cognitive and more generally somatic ageing does not appear to be a strict function of chronological age in eusocial insects, and the social role of the animal into the hive seems to act as a modulator of biological ageing [40].

While data on ageing in eusocial insects reveals intriguing caste-specific patterns, investigating the ins and outs occurring specifically in long-lived queens remains a challenge. One potential factor contributing to their extended longevity is the shielding of queen physiology induced by reproduction *per se*. It has been observed that mating can actually increase the longevity of queen ants *Cardiocondyla obscurior,* regardless of whether the males have been sterilized or not [43]. Moreover, unlike reproductively active queens, non-reproductive queens of this species (identified in the study as not laying eggs at all) exhibit a comparable or even lower life expectancy than workers when they are kept solitarily [44]. Interestingly, studies involving bees and ants have shown that when the queen is removed, and workers start reproducing, they tend to live longer than their counterparts which do not reproduce in colonies with a queen [45–47]. Furthermore, in queenless species such as *Diacamma sp.,* all females have a spermatheca and can theoretically mate at any time, but those that actually start being breeders (i.e., reproductive workers also called gamergate) also live longer than non-reproductive individuals even if they are indistinguishable from each other in terms of development and morphology. In laboratory, *Diacamma sp.* and *D. cyaneiventre* show a similar trend with average adult lifespan being three times longer for gamergates [48,49]. Here, neither mating nor insemination appears to extend gamergate life span, as non-mated workers that become reproductive also live longer. One hypothetical explanation for this paradoxical longer life expectancy of breeders is that workers assume a higher workload, a general pattern in social insects [49]. Negroni *et al* have investigated the transcriptomic regulation of lifespan extension in ant fertile workers of *Temnothorax rugatulus* [47]. They found that the most differentially expressed gene, encoding an isoform of catalase, was downregulated in reproductive individuals. Long-lived reproducers might therefore be exposed to lower levels of oxidative stress [47]. This situation is mirrored in the sub-family of ponerine and or in honeybees, where antioxidant enzymatic activity is lower in gamergates or queens than in workers [50]. On the other hand, there is an increase in catalase transcript abundance and activity in termite queens [51], and the same observation is made for Cu/Zn-superoxide dismutase (SOD) activity [52]. It is noteworthy that catalase is also upregulated in fat bodies of old *versus* young queens of *Temnothorax rugatulus*, suggesting a late-life up-regulation of anti-oxidant shielding that may regulate ageing rate [53]. Thus, it seems that taxon-specific (ants *vs.* termites) patterns of antioxidant mechanisms may have differently evolved in the regulation of caste-specific longevity among social insects [54].

The mitochondrial function among ant castes of contrasted longevities is likely to bring key additional data to the so-far above story. Since all females in a given colony inherit their mitochondria from the founding queen, we sought to explore how mitochondria from a unique maternal line are actually associated with different longevities. We assessed individual metabolic rate (oxygen consumption), mitochondrial density and functioning (efficiency in ATP production, energy charge), as well as different proxies of oxidative status in queens and workers of *Lasius niger*. In line with the predictions of the oxidative stress theory of ageing, we hypothesized that mitochondrial characteristics should differ among castes, with long-lived queens more likely to exhibit a bioenergetic profile associated to long lifespan, *i.e.* low ROS production, high antioxidant capacity and therefore less cellular damage for a given level of oxygen consumption and/or ATP production.

## Material and methods

### Ant segregation/ caste identification

The detailed methodology of *Lasius niger* segregation is provided in the supplementary methods, Supplementary Material. Briefly, queens are easily identifiable due to their larger size, and workers were classified based on their social roles: nest-workers cared for the queens and eggs, while foragers searched for food outside the nest. We assume foragers are mostly older due to age polyethism in their division of labour [55]. Workers were pooled in groups of 5-7 individuals for indirect calorimetry (some escaped during the initial protocol step just before oxygen consumption measurements). The same ants were then used for different enzymatic activities: citrate synthase, aconitase, fumarase, and glutathione, respectively. Other ants were segregated in the same way and employed for independent measurements. Sample sizes for each experiment are summarized in Supplementary Table 1. In total, 28 queens and 195 workers were used.

### Oxygen consumption (indirect calorimetry)

We measured O2 consumption from foragers, nest-workers and queens using an Oxygraph O2k (Oroboros Instrument GmbH, Innsbruck, Austria) and DatLab software. The system is normally used with stirred liquid suspensions such as isolated mitochondria. To proceed with a gaseous atmosphere, we followed the protocol detailed in [56]. The 6 queens (4 years-old) and the 22 worker pools for which the metabolic rate has been measured were done over 2 sessions.

### Body mass and metabolic rate calculation

All pools were weighed directly after oxygen measurements (Mettler Toledo AG285, ± 0,01mg accuracy). To improve comparison between measurements made at different days, two calibration weights of 100 mg and 10 mg were prepared with platin wire and used as references. Mean body mass values in mg were (mean ± SD): queens 28.19 ± 5,419, foragers pools 4.271± 0.5287, nest-workers pools 5.147± 1.055. We calculated the metabolic rate by dividing the oxygen consumption rate by the pool mass raised to the power of a caste-specific allometric coefficient (α), as previously determined [56, under review]. After having their body mass measured, the ants were placed in a 15 mL falcon tube with cotton impregnated with water before being ground for subsequent measurements of enzymatic (aconitase, fumarase and citrate synthase) and glutathione content.

### Preparation for enzymatic measurements

The very next day of metabolic rate measurement sessions, ants were mashed into 300 µL phosphate buffer (0.1M, 5mM EDTA, triton 0.1%, pH 7.3). Homogenates were immediately placed in dry ice and defrosted at room temperature, and centrifuged at 8,000 g at 4°C for 10 min. Resulting supernatants were used for enzymatic measurements. Aconitase/fumarase activities were assayed within 2 hours, other tubes were aliquoted and kept at -80°C for the measurement of the citrate synthase activity.

Homogenates from *Locusta migratoria* flight muscles was previously frozen at -80°C and used as an inter-plate controls. Normalization of enzymatic activities was done using samples’ protein content determined by BCA assay (Pierce Company).

### Glutathione content

Total glutathione (GSH) content was determined by the glutathione reductase enzymatic method [57]. We used half area plates to concentrate samples (Greiner Bio-One 675101) with small volumes of reagents. Glutathione standards were prepared from a 3.28 mM GSSG stock solution for final concentrations from 0.048 to 3.78 μM. A working solution of 5 U/mL of glutathione reductase and 2.6 mM of 5,5′-dithiobis(2-nitrobenzoic acid) (DTNB) was prepared in assay buffer. All samples were deposited in duplicates. Reaction was initiated by mixing 20 μL of the latter working solution with 35 μL of homogenate/ standard. We used 1:12 and 1:3 sample dilution for queens and workers respectively. Finally, 10 μL of 1.2 mM NADPH solution were added and the increase in absorbance was recorded at 405 nm wavelength (final volume of 65µL), with a TECAN spectrophotometer (model Infinite M200). Glutathione concentrations were normalized again using sample protein content (see above).

### Aconitase/fumarase activity ratio

The aconitase and fumarase activities were measured in 96 well plates allowing measurement at 240 nm wavelength (Greiner UV-Star® half area 675801) with the TECAN spectrophotometer. The test is measuring the rate of activity (slopes) of those enzymes in presence of increasing substrate concentrations. Firstly 50 µL of Phosphate buffer (100 mM phosphate buffer, EDTA 5mM) were introduced in each well, followed by 20 µL of the homogenate / 20µL buffer for negative control (1:4 dilution for Queens because optical density (OD) was too high *per se*). Samples were run in duplicate. The reading protocol for measurement of 240 nm wavelength was as follows: (1) Shaking (15 s, 2mm linear); (2) readings every 20s on 2min to evaluate the background and stability; (3) aconitase reaction with addition of isocitrate solution 120 mM (25 µL) with the aid of a multichannel pipette, shaking as before, 8 minutes of readings each 20s with 3s shaking (slope a); (4) fumarase reaction with addition of malate 250 mM (25µL), rest of the procedure as before (slope b). Aconitase to fumarase ratio was calculated with the assumption that increase in OD summed the rates of aconitase and fumarase reactions (see supplementary Figure S1A). Hence, slope for fumarase was calculated as slope (a) – α slope (b), with α taking into account the dilution induced by next reagents (α =19/24 for theses volumes). Activities were calculated with ε_ac_ = 3.410 and ε_fum_ = 2.440 M^-1^.cm^-1^ for aconitase and fumarase respectively before expressing the ratio [58,59]. The activity was calculated as follow:

Activity : V_sample_/V_reaction_ * 60 *(slope/(ε*L)

with L the optical path length (cm) depending on the reaction volume of the half area plates (0.588 and 0.730 cm). Aconitase and fumarase activities were normalized by protein content (BCA assay), and we verified aconitase sensitivity to H_2_O_2_ in our animal model (Figure S1B).

### Citrate synthase activity

Citrate synthase is a Krebs cycle enzyme (in the mitochondrial matrix) and its activity is generally used as a proxy for mitochondrial density/activity. Citrate synthase activity (CS) was carried out at room temperature (25°C) using the TECAN spectrophotometer by measuring the appearance of the CoA-SH [acetyl-CoA + oxaloacetate ↔ citrate + CoA-SH + H^+^ (side reaction: CoA-SH + DTNB → yellow TNB)]. For the CS assay, and as suggested by Eyer and collaborators [60], the molar extinction coefficient used in a phosphate buffer 0,1M at 25°C was ε=14 150 M^−1^ cm^−1^ for DTNB at 412 nm wavelength. The reaction mix consisted of 135µL of phosphate buffer, EDTA 5mM, DTNB 0.11mM, and 0.33 mM acetyl CoA to which 5µL of sample homogenate were added. Reaction started with addition of 10µL of 10mM oxaloacetate *per* well (150 final volume). Kinetics were conducted during 5 minutes with 15 seconds interval measurements. Samples were deposited in duplicate into half area plates, and results normalized by sample protein content.

### Catalase activity

The catalase measurements were taken from pools of ants segregated in parallel with the previous experiments, but the colonies used here were young-of-the-year (so were the queens). The enzyme activity was obtained with the Oroboros reading, based on the following reaction: H_2_O_2_ -> H_2_O + ½ O_2_. Calibration was done in water, chambers opened for 100% oxygen reference, closed with dithionite addition for 0% oxygen reference. A single segregated ant was mashed into 530 µL phosphate buffer (0.1M triton 0.1%, 5mM EDTA, pH 8) and placed into 0.5 mL chamber volume. We used a data recording interval of 2 seconds with background correction, stirrer 750 rpm at 25°C (Figure S2). Slope was calculated on 10 points after injection. We injected 2µL H_2_O_2_ into the chambers (final concentration 500µM, H_2_O_2_concentration checked at OD 240 mm). At the end of acquisition, each mashed sample was collected for normalization by protein content (BCA assay). We also injected H_2_O_2_ into chambers containing phosphate buffer to verify that oxygen production was only due to biological content.

### Mitochondrial density by qPCR and electron microscopy

Mitochondrial density was first measured using qPCR amplification method. DNA extraction was done with TRIzol^TM^ following the procedural guidelines, with volumes 1:2, that means ants were crushed into 500µL of TRIzol^TM^ on ice (3-4 mg fresh material for worker pools). The homogenization step was followed by a first centrifugation at 8000g, 3min and 4°C after 5 min incubation, to remove cuticular debris. Samples were resuspended in 40 μL of DEPC-treated water at the end of DNA purification. The single-copy gene long wavelength rhodopsinpreviously (LW-Rh) defined by Quque [61] was used as reference. The primers’ sequences were: LW-Rh-Forward 5’ - GGA CCC TTG TTC TGT GAA CTG TA - 3’, LW-Rh-Reverse 5’ - ATT ACG TTG TAC CTG TCG AAT GC - 3’ for reference, Ln-COI-F 5’ – ACC TGA TAT AGC ATA CCC CCG T -3’, Ln-COI-R 5’ – AAC AGT TCA TCC TGT TCC TAC TCC -3’ for mitochondrial COI gene and Ln-16S-F1 5’ – CCG CAG TAT TTT GAC TGT GC -3’ Ln-16S-R2 5’ – TCC TTC ATT CCA GTT CTT AAT -3’ for mitochondrial 16S gene.

Mitochondrial density of brain tissue was also assessed using electron microscopy. One ant of each group was used (one queen and one ant for each workers’ subcastes). See supplementary methods for complete details. Mitochondrial parameters such as densities, size, volume and shape (circularity index which indicate fission/fusion propensity) were measured on Fiji software following Lam protocol [62]. The higher the circularity index, the less frequent mitochondrial fission/fusion events [62]. Multiple image captures were taken under the microscope from a single preparation (4 sections), enabling us to perform statistical analysis on the results. Unfortunately, the brain sections from the nest-worker did not seem to originate from the same region, so we set them aside. Counting of lipofuscin aggregates were also conducted on these image captures (see supplementary).

### Adenylate and guanylate energy charge and ATP content

The Adenylate Energy Charge (AEC), proposed by Atkinson in 1967 [63], is used as a practical index of the physiological status and health of the cells. It is given by the following formula: AEC = ([ATP] +0.5[ADP])/([ATP] + [ADP] + [AMP]). The normal AEC for viable organisms (eukaryotes as prokaryotes) is comprised between 0.8 and 0.9 [64,65], i.e. ATP producing reactions are in balance with the ATP consuming reactions. When AEC drops of between 0.7-0.5, organisms are still in a viable state, but homeostasis is not maintained [66]. Very unfavourable conditions are when AEC<0.5, a status lethal for most cells [67]. Guanylate Energy Charge (GEC), an analogous to AEC but for guanine nucleotide, is also an indicator for metabolism, that reflects the cell ability to synthesize proteins [68]. The detailed procedure is provided in the supplementary methods.

### Principal Component Analysis of proteomic mitochondrial content

We conducted an *in silico* reanalysis of proteomic data previously acquired by our group on ant colonies of the same origin [69]. On the 1,380 proteins previously quantified, 240 proteins were identified as playing a role in the mitochondrial function. Data analysis with DEP v1.22.0 package was conducted focusing on those mitochondrial proteins’ expression among queens, nest-workers and foragers on the basis of fold changes (calculated based on Limma V3.56.2) as developed in [70]. To properly run the analysis, missing values that concerned all individuals in a same group, or even a given caste (see supplementary Figure S3A), were assumed as missing not at random (MNAR) and were attributed the minimal quantitative value in the dataset; other missing data were assumed as missing at random (MAR) and inferred with the *k*NN (k nearest neighbours) algorithm (DEP uses VIM package v6.2.), which imputes missing values based on proteins with similar expression profiles [71]. A PCA was executed on the mitochondrial dataset to visualise the contributions of castes and proteins to the orthogonal independent axes (Figure S3B). Fold changes were illustrated using volcano plots (Figure S4), and statistically significant results are given in Table S2.

### Statistics

Statistics analysis were conducted using RStudio v4.0.2 (http://www.R-project.org/) and prism v9.4.0. Normality of model residuals was verified by Shapiro-Wilk test, homoscedasticity by Bartlett’s test. For each random variable, social group (queen, nest-worker or forager) was used as fixed factor. One way ANOVA with Tukey’s multiple comparisons test were conducted when normality and homoscedasticity were verified (O_2_ consumption, O_2_ consumption reported to the mass, AEC, ATP/ADP, ATP content, GEC and aconitase/fumarase ratio). In case of unequal variances, Welch ANOVA test was employed with Dunett’s T3 to compare all pairs of variables (mass, citrate synthase activity, ATP/AMP ratio, fumarase activity, aconitase, fumarase and catalase activities). In conditions where normality has not been verified, comparison on ranks was done with Kruskal-Wallis test (inosine, inosine reported to the mass and glutathione levels). Similarly, paired comparisons were done when workers were merged or a subcaste absent, with Welch’s t test in case of lack of homoscedasticity (for mitochondrial volume and density), otherwise Mann-Whitney tests were used for non-parametric data (qPCR analysis and circular index). Specific post-hoc test P-values are indicated on the figures.

## Results

### Queens have the lowest metabolic rate, old nest-workers the highest

Body mass between the three social castes were significantly different (F_(2, 20.30)_ =165.4, P < 0.0001). Queens were 32 times heavier than nest-workers, and 39 times heavier than foragers (Supplementary Figure S5A; P < 0.0001). The weight of nest workers was also significantly 22% higher than that of foragers (P < 0.05).

Long-lived queens had half the metabolic rate of workers (F_(2, 29)_ = 75.87, P <0.001) and a higher metabolic rate was found in foragers compared to nest-workers (adjusted P = 0.0431) due to their weight, absolute oxygen consumption between the workers being not different Figure S5B (no significant differences, adjusted P = 0.9584).

### Markers for mitochondrial abundance and ATP production

Citrate synthase enzymatic activity (Figure S6A) was more than 6 times higher in workers than in queens (F_(2, 10.49)_ =62.08, P <0.0001), with no difference between the two groups of workers (P = 0.9637). For mitochondrial density estimated by qPCR, a median value of 122.5 was obtained for queens versus 295.5 (twice higher) for workers (Figure S6B). However, queens could not be significantly discriminated from workers (foragers and nest-workers not analysed separately, Mann–Whitney *U* = 14, P = 0.3660 two-tailed).

Lastly, different mitochondrial parameters such as density and volume in the brain of queens and foragers were assessed using electron microscopy (Figure S7). Unpaired t-tests with Welch’s correction showed no difference in mitochondrial volume (which occupy between 6.5 and 7% of the cell volume), nor in mitochondrial density (t=0.4806, df=4.317 and t=1.305, df=4.364, respectively). Interestingly, the assessment of organelle shape after outlining highlighted a significant difference in the circularity index (index median of 0.5355 and 0.8260, in foragers and queens, respectively, Figure S7C, P < 0.001), thereby indicating that fission/fusion was more common in queens than in foragers.

As a complement to mitochondrial density proxies, the PCA analysis on proteomic data on mitochondrial proteins from Quque *et al* [69] was performed (Figure S3). The first two axes explained 86.9% of the variance with the first axis differentiating queens (positive PCA1 values) from workers (negative PCA1 values), while PCA2 less clearly differentiated workers among them. Based on proteins loading on PCA axes, the abundance of only 22 proteins was significantly different among groups for a P < 0.05 (supplementary Table S2): 17 were differentially expressed between queens and both workers castes, 9 between queens and nest-workers, and finally 5 between worker castes (see Table S1 and discussion for details).

Mitochondrial ATP content was not statistically different in the three groups (F_(2, 6)_ = 1.753, P = 0.2514, Figure S8A), despite a trend towards higher ATP content in foragers.

Finally the AEC index (Figure 1B) was significantly higher in the queen caste compared to workers: queens mean value was of 0.934, while it was of 0.898 for nest-workers and of 0.888 for foragers (F_(2, 20)_ = 13.80, P < 0,005). All AEC values recorded here were > 0.8, indicating that none of the individuals were in a marked energy stress status. Consistent with the AEC from which it is derived, ATP/ADP ratio (Figure S8B) was 55-70% higher in the queen caste compared to both worker castes (ratio of 9.48, F_(2, 20)_ = 12.02, P < 0.005). Likewise, ATP/AMP is two to three times higher in queens Figure S8C (F_(2, 9.95)_ = 8.712, with P < 0.005).

**Figure 1:**
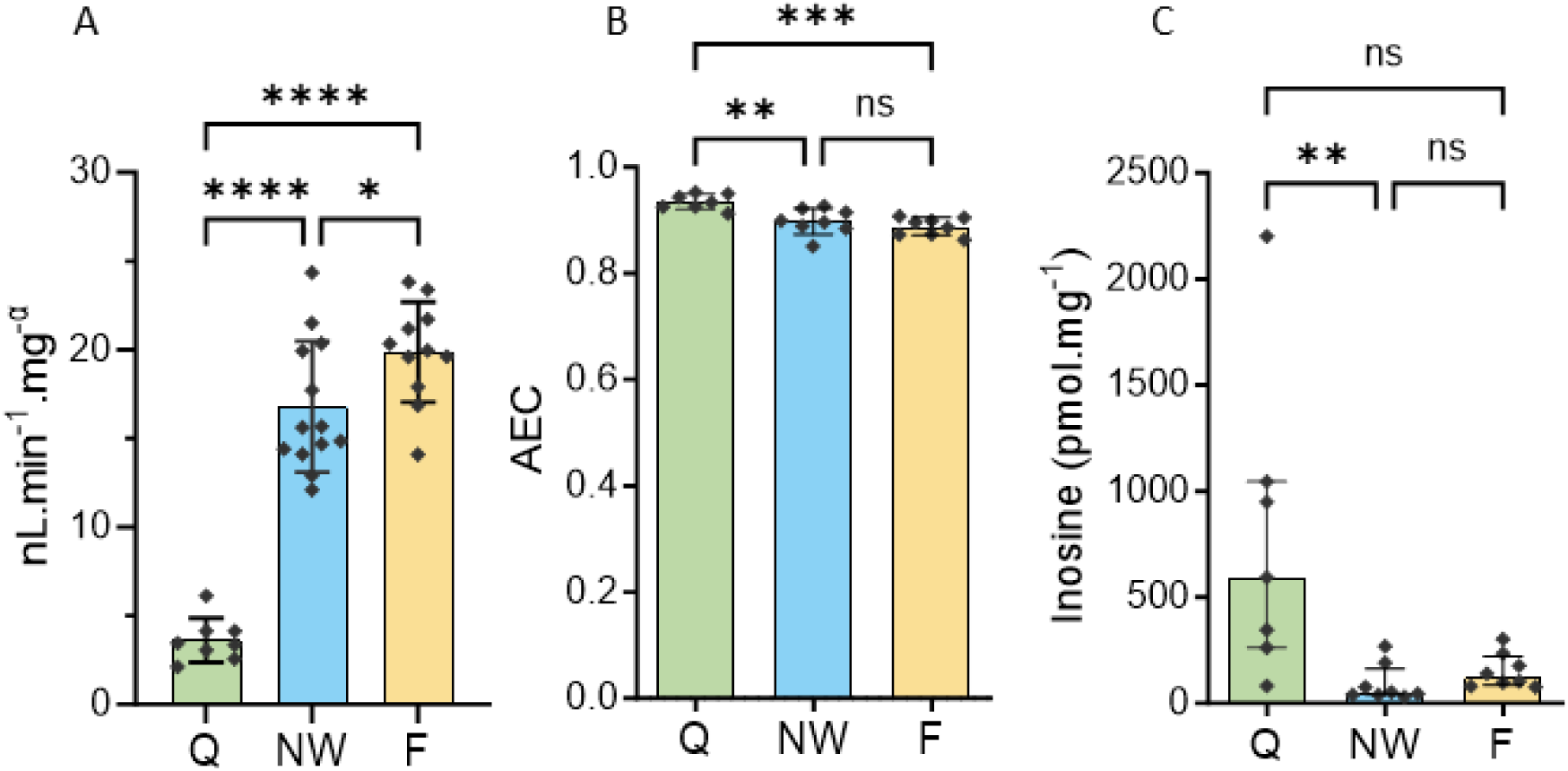
Metabolic rate (A), with α as the allometric coefficient, adenylate energy charge (AEC in B), and inosine normalized by the mean mass of each caste (C). Values are mean ± SD in A and B, median (IQR) in C. Q: queen, NW: nest-worker, F: forager. ****P <0.0001, P**<0.005 P*<0.05.

There were no significant differences in Guanylic acid Energy Charge (GEC) between castes (F_(2, 20)_ = 0.5760, Figure S9A). Nonetheless, we observed consistent peaks of inosine (component of the salvage purine synthesis pathway) in the queen caste (Figure S9B). As individuals were not weighed during the experiments to avoid any stress, inosine was expressed in relation to the mean mass obtained from the entire population (Figure 1C). The level of queen inosine was significantly higher in queens than in nest-workers (by a factor of 10, Kruskal-Wallis test H = 11.62, adjusted P< 0.005). Inosine level was also five time higher in nest-workers than in the forager group, but statistical significance was not reached (adjusted P = 0.1435, 0.4211 between workers).

### Worker brains show lipofuscin deposits, with higher catalase activity

The activities of aconitase and fumarase were found significantly higher only in nest-workers compared to the queens (F_(2, 11.80)_ = 8.938, P <0.01; F_(2, 10.98)_ = 8.303, with P < 0.01, supplementary Figure S10 A and B, respectively). The aconitase/fumarase ratio was not different among the three castes (F_(2, 21)_ =0.1037, with P = 0.9020, Figure 2A).

**Figure 2:**
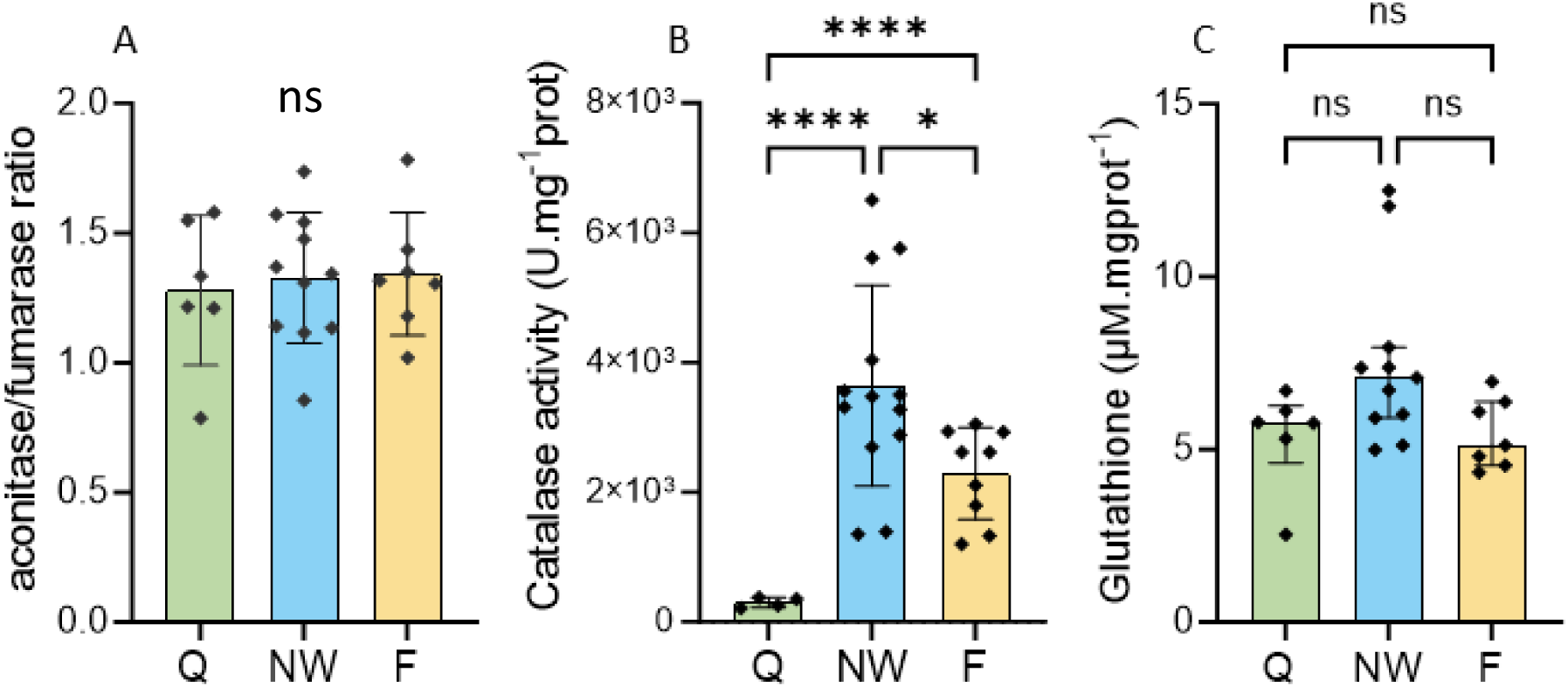
The aconitase/fumarase ratio as indicator of oxidative stress (A), and the antioxidant capacities measured via the activity of catalase (B), and glutathione concentration (C). Values are mean ± SD in A and B, median (IQR) in B. Q: queen, NW: nest-worker, F: forager. ****P <0.0001, P*<0.05.

In addition to enzymatic measurements, electron microscopy allowed us to observe lipofuscin deposits highlighting lamellae structures (fingerprint) in ants’ brains as shown in Figure S10C. The sample size was too small (n=1, there was no lipofuscin on all sections) to conduct proper statistical analysis, but many synapses had lipofuscin in the foragers’ brain (up to three distinct structures in a same synapsis), and two structures were found in the nest-worker sections while none were characterized in the brains of queens (which were older than the workers).

Catalase activity, shown in Figure 2B, was highest in the nest-worker group (F_(2, 13.26)_ = 60.56, all P < 0,0001) than in the queen (12 times lower) and the forager castes (6 times lower). No significant difference was found between worker castes, P = 0.0370). No statistical differences in glutathione measurements were found between castes in Figure 2C (Kruskal-Wallis test H=6.105, P <0.05).

## Discussion

Our comparison of mitochondrial functioning among *Lasius niger* castes underlined several key bioenergetics characteristics. First, measurements of oxygen consumption rate and Krebs cycle enzymatic activities (citrate synthase, aconitase and fumarase) indicate a lower mitochondrial activity in queens compared to workers. Second, the mitochondrial volume and mtDNA/nucDNA ratio suggested that the mitochondrial content is similar in queens and workers. Third, the energy charge of queens’ mitochondria was higher. Fourth, oxidative stress appeared to be of lower amplitude in queens with (i) for a comparable level of aconitase/fumarase ratio, (ii) a stable (glutathione) and lower (catalase) antioxidant defence levels, (ii) proteomic profiles suggesting a specific antioxidant protection of key mitochondrial enzymes or enhanced mitochondrial fission/fusion rates (i.e, a proxy of mitochondrial maintenance) and (iii) less (no detection) of lipofuscin. Overall, our data suggest that ant queens may live longer because they combine lower oxygen consumption with higher ability to ensure mitochondria maintenance, but not from increased antioxidant defences.

The measurement of adenine nucleotides highlighted that AEC was higher in queens. ATP hydrolysis is the primary means of supplying energy to processes that prevent fitness decay in the short term (e.g., preservation of cellular ions gradients, active elimination of waste/incoming poisons) and in the long term (e.g., replacement of altered/aged structures needing protein and lipid synthesis, DNA repair). Adenosine triphosphatases (that convert ATP into ADP, hereafter ATPases) are sensitive to the concentration of their substrate (ATP) and to the ratio between substrate and product (ATP/ADP ratio). Moreover, the higher the ATP/ADP ratio, the greater the energy released by ATP hydrolysis [72]. Consequently, with higher ATP/ADP ratio, ATP consumption effectors (ATPases) in queens are beneficiating from a more efficient use of energy than those in workers. Such a situation would particularly favour ATPases when the cell is short in energy, like those involved in protein recycling including mitochondrial proteins [73], and would guarantee better somatic protection/repair of the queen’s organism as a whole. Replacement of non/moderately damaged biological structures is, in the short term, a futile cycle but in the long-term impacts positively fitness by mitigating damage accumulation (ageing). The extreme difference in longevity observed between closely related individuals (queens and worker ants) was expected to reveal previously described biochemical factors associated to the fitness of the organism (e.g., oxidative stress), not directly under the control of genes (as ants of a same nest are close kins) but rather of lifestyle (determined by castes). Our results suggest that an higher AEC could belong to these factors, as its decrease would mechanically reduce global ATP availability but also the absolute hydrolysis energy gain by molecule of ATP (due to the Gibbs free energy effect) [72,74]. Again, the ultimate consequence of such bioenergetic advantage will be reflected in a higher rate of renewal of biological structures and a reduction of damages: in biochemical terms, queens are therefore permanently younger. Workers’ lifestyle induces a higher energy demand, *i.e.*, a higher ATP turnover rate and consequently a higher metabolic rate. As a consequence, for ATP production to match the energy demand, the metabolic rate will rise and/or the AEC will decrease (with an increase in AMP, the lowest energy state of adenylate). As a consequence, the high ATP consumption that sustains worker lifestyle may have a long-term deleterious impact on the maintenance of the soma, by reduction of the renewal rate of biochemical structures (including mitochondria themselves). Any unbalanced investment in mitochondrial maintenance will inevitably lead to a higher fraction of damaged components and a faster decay of performance of workers (senescence), ultimately resulting in a shorter lifespan. Whether such an energy ATP-dependent trade-off mechanism is effectively modulating the levels of oxidative damage deserves consideration, as it would explain how ageing is dictated by the social status (worker or queen) in eusocial insects. Studies carried out on isolated mitochondria do not show that ROS are produced as a direct consequence of increased oxygen consumption, but rather the opposite [3]. In contrast, altered enzymatic function of redox proteins (of the electron transport chain) tends to favour the release of mitochondrial ROS thus implying a higher demand in terms of repair, a situation at risk on the long term due to the progressive decline of repair performance with age. In the following discussion, we will highlight that queen ants may actually have coevolved a bioenergetic status compatible with a higher investment in mitochondrial maintenance.

Black garden ant queens were found to have a lower (based on citrate synthase activity), or at least comparable (based on mitochondrial genome copy numbers or on total mitochondrial volume) mitochondrial density to that of workers. In the literature, mitochondrial density is often described to increase with advancing age to compensate for altered mitochondrial function due to oxidative damage accumulation, for instance as stated in the trophocytes and fat cells of worker bees [75]. Previous observations of young and older queen bees (16 months) found no difference in mitochondrial density [76], suggesting that the maintenance of their bioenergetic capacity with age is a particular trait of this long-lived caste. The higher AEC index in ant queens supports the idea that they have a higher capacity of energy production per mitochondria compared to workers. Additionally, they have significantly higher amounts of inosine, which could suggest increased utilization of the purine salvage pathway in the biosynthesis of corresponding nucleotides, thereby avoiding the higher energy costs of *de novo* purine biosynthesis for their cellular functioning [77,78].

Proteomic profiles in ants highlighted higher abundances for a few proteins in queens, notably proteins of the mitochondrial membrane (see summary diagram Figure S11). Voltage-dependent anion channel 1 (VDAC1) is the most abundant protein in the mitochondrial outer membrane [79], and the two porines we identified were both more abundant (up to 14-fold) in queens. VDAC1 is important for a variety of mitochondrial functions, including diffusion of metabolites (such as ATP, ADP, NADH, pyruvate), nucleotides and ions (such as Ca^2+^) at the outer mitochondrial membrane [79]. The higher expression of VDAC1 supports the idea that ATP/ADP traffic is a key parameter of queen bioenergetics. The most significant difference between queens and workers concerns oxoglutarate dehydrogenase (OGDH, or 2-oxoglutarate dehydrogenase complex component E1, also referred as KGDH for α*-* ketoglutarate dehydrogenase). A higher content of associated 2-oxoglutarate malate carrier (OGC) that transports the 2-oxoglutarate (or α-ketoglutarate αKG) protein across the inner membranes of mitochondria is in line with the idea of a higher mitochondrial maintenance process. OGDH has been shown to maintain mitochondrial fusion and fission events in both *C. elegans* and human cells [80]. These proteomic data support our indication of more frequent mitochondrial fission/fusion events also suggested from electron microscopy and the higher circulary index measured in queens (see Figure S7C). As altered mitochondrial dynamics with senescence can inhibit mitophagy and lead to accumulation of damaged or dysfunctional mitochondria in cells [81], higher rate of fission/fusion events, by regularly rejuvenating the mitochondrial pool, should better protect queens from the deleterious effects of ageing on bioenergetics.

A high metabolic rate and oxygen consumption implies an inevitable associated risk of oxidative damage to biomolecules by ROS. The ’ROS theory of ageing’, which complements the rate of living theory, stipulates that the higher the metabolic rate and the associated oxidative cost, the faster the ageing process. Contrary to our initial expectations, the aconitase/fumarase ratio was not found to be higher in the long-lived and less metabolically active queen caste compared to the worker caste. Still, even if this observation needs confirmation, lipofuscin, another proxy of ageing, was found at higher levels in the worker microscopic sections (supplementary Figure S10C). This auto-fluorescent pigment has been observed to accumulate in other insects like social bees with chronological age [82,83]. Since the inhibition of mitochondrial fission leads to increased lipofuscin formation [83], which might hinder autophagy and disrupted mitochondrial degradation, the lower lipofuscin signal in queen reinforces our previous hypothesis that mitochondrial dynamic is more prevalent in queens and/or results in a faster accumulation of dysfunctional mitochondria as worker ants age [75].

The antioxidant barrier was found to be lowered in the long-lived queens, as our measurements of catalase enzyme found it less active in queens (or non-significantly different for glutathione levels). Thus, the antioxidant parameters are not correlated with the long-lived phenotype in the black garden ant, contrary to other social insects such as termites [51,52]. It may be that queens do not need to invest in antioxidant compounds as they may benefit from (i) a lower rate of mitochondrial ROS production, (ii) the social shielding as workers are the only ones on the front line of environmental stress (e.g., [84]). In certain ant species, intense cooperation relies on trophallaxis, the mouth-to-mouth exchange of fluids within the colony, and the division of labour results in a chain of food dissemination in the form of a hierarchical network [85,86]. Proteomic analysis of trophallactic fluid in colonies of the ant *Camponotus floridanus* revealed that nurses, responsible for brood care and closer to the queen in the social network, have higher levels of proteins involved in oxidative stress response (superoxide dismutase and glutathione peroxidase). Trophallaxie might therefore in this case help the individuals at the end of the food chain to benefit from the costs that others individuals belonging to the same social network actually incur [87].

In conclusion, our work supports a mitochondrial basis for the unexpected longevity of the black-garden ant queen. This process extends beyond the simple ROS/antioxidant equation, as we show that the queen’s longevity appears based on (i) a higher ATP/ADP ratio leading to a more efficient energy release for each molecule of ATP hydrolysed, (ii) a lower metabolic rate and (iii) a higher investment of saved energy in the maintenance of mitochondrial functioning as reflected by the regulation of fusion/fission or the expression of mitochondrial enzymes of the Krebs cycle. The enhanced maintenance of mitochondria may be a key component of the queen’s particular metabolism and the paradoxical trade-off between the high metabolic demand of reproduction, their relatively lower metabolism and their exceptional longevity.

## Ethics

The colonies used in this study came from wild, newly-mated queens captured by us, or purchased from a specialty store. Import and export licences are not required for the transport of our study species. No ethical approval was required from an animal welfare committee.

## Authors’ contributions

MK, FB and FC set up the experiment, MK carried out the measurements and statistical analyses, MK and FC wrote a first version of the manuscript and all co-authors drafted the final manuscript and agreed to publication.

## Acknowledgements

We acknowledge the Platform facility of Cochin Institute for electron microscopy, and give all thanks to SAM Platform for its prompt service and quick responses to our questions. The Integrion chromatography station was acquired with the financial support of SIRIC BRIO2 (COMMUCAN) and Region Nouvelle-Aquitaine (AAPPF2021-2020-12000110). The authors express gratitude to Mary Lucienne Waller for proofreading the language.

## Funding

The ETERNEEL project was funded by a grant (2020-2021) from the MITI agency (Mission pour les Initiatives Transverses et Interdisciplinaires) of the CNRS (Centre National de la Recherche Scientifique), who also granted M. Kervella with a PhD grant (2020-2023).

## Conflict of interest declaration

We declare we have no competing interests.

## Notes

### Competing Interest Statement

The authors have declared no competing interest.

