## Supplementary methods for "Mitochondrial maintenance is involved in the exceptional longevity of reproductive queens of the eusocial ant *Lasius niger*"

Ant segregation

The majority of the colonies used in this study were derived from wild, newly-mated *Lasius niger* queens (Linnaeus, 1758), which were captured at the “Campus Plaine” site of the Université Libre de Bruxelles (50°49'08.4"N 4°23'57.0"E). These queens were reared in the laboratory for four years, initially at [name of the lab], France, for the first three years, and then relocated to Paris for measurements in the final year. The youngest colonies were ordered online through the website antstore.net and were from the current year.

Colonies were maintained at room temperature (25°C) with a relative humidity of 50-60% during the season, and were placed in diapause from December to February at 10°C (gradual decrease in temperatures). A test tube served as the nest, with water at the bottom of the tube separated by a cotton barrier to maintain humidity. Each tube was placed inside a plastic container measuring 20 to 30 cm on each side, with one nest per container due to inter-colony aggression (Fluon applied along the edges to prevent ants from escaping). The colony sizes varied, ranging from about twenty to sixty workers for a queen (monogyny species), depending on their age. The ants were provided with a diet of 0.3M sugar water and defrosted mealworms once a week.

In *L. niger*, there is an age-related division of labour [1], the youngest workers taking care of larvae (in-nest task, called nest-workers thereafter), while the oldest ones are devoted to out-nest tasks (as foraging [2], called foragers thereafter). Since there is no dimorphism between them, we separated workers according to their behaviour, as described in [3]. Briefly, the foraging behaviour was stimulated by a 4-day fast, at the end of which a high concentrated sugar solution (0.3 M) was supplied in a plastic tray. Individuals repeatedly found active in the foraging area were identified as foragers, whereas the ones repeatedly found cloistered inside the nest were identified as nest-workers. The tube used as anthill was therefore clogged with cotton for trapping foragers outside and taking the time to mark their abdomen with an acrylic ink (Posca®). Ants were anesthetized by cold if necessary (0.5–1 min, on ice). We then collected the nest-workers and marked them with another colour. We particularly spotted those showing tendency to form immobile clusters as it has been described in *Caponotus fellah* [4]. All ants were put back into their colony where usual food and water were provided for 2 days. The fasting process was repeated 3 times, with alternating colours to avoid marking nest-workers which were too curious due to food deprivation. The ants finished being segregated a week before the start of the experiments (48 hours fasting).

Measurements were made between November and December. Since tagging was repeated, we assumed for our study that there was not any switch in ants’ behaviour, and that foragers were the oldest workers, without knowing their exact age. Actually, we rarely observed starved nest-workers going out in search of food, and we almost exclusively marked the same ants. As we did not either remove the eggs progressively we can assume more heterogeneity among nest-workers.

Electron microscopy

Ants were individually anesthetized on ice, and immediately dissected into fixative under binocular magnifier (mitochondria are first elements altered by ambient air), after needles were placed under the head and into clypeus (from the top before propharyngeal gland) to facilitate an opening of the head. Brain was recovered. The fixative was composed of glutaraldehyde diluted to 2.5% in phosphate-buffered formaldehyde solution (ROTI®Histofix 4 %), pH adjusted to 7.2 with a few drops of KOH. Brains were washed in phosphate buffer, post-fixed in 2% osmium tetroxide, dehydrated in graded ethanol, and embedded in epoxy resin. Ultrathin sections (50–90 nm) were cut on an ultramicrotome (8800 Ultrotome III; LKB Bromma) and collected on 300-mesh nickel grids. Staining was performed on drops of 4% aqueous uranyl acetate, followed by Reynolds lead citrate. Ultrastructural analyses were performed with a JEOL JEM-1011 electron microscope and digitalized with the DigitalMicrograph software.

Adenylate/guanylate energy charge and ATP content

Ants have been segregated more than a week before measurements (since any stress can decrease AEC value). They were caught gingerly and directly ground with Dounce homogenizer in an ethanol/Hepes 20 mM (7/3; v/v) solution -precooled at -20°C of. Glassware and solutions were kept on dry-ice. Samples were stored at -80°C a few days before being sent to SAM facility (TBMCore Bordeaux). Ethanol/Hepes extracts were evaporated using a rotavapor (3min, 65°C) and the dried residue was resuspended in 500μl of nuclease/phosphatase-free water. Insoluble particles were removed by centrifugation (21,000 g, 4°C, 1 hour) and the supernatant was ultra-filtrated on nanosep10K Omega (Pall). Metabolites were then separated on an Integrion chromatography station (Thermo Electron) at 0.38 ml/min and 30°C using an AS11-HC-4μm column (250 × 2 mm, Thermo Electron) and the potassium hydroxide discontinuous gradient as described in [69]. Peak area quantification was done by UV absorbance at 260nm for all nucleotides/nucleoside. Metabolite contents were determined using standard curves obtained with pure compounds.

ATP content was also measured on 3 other whole ants per group, with ATP Bioluminescence Assay Kit CLS II by Roche, Mannheim Germany. Since the pH optimum is between 7.6 and 8, we increased tricine concentration as suggested in the protocol (from 0.04 to 0.1M here) to avoid our samples acidification due to formic acid. The pH never dropped off even before dilution preceding luminescence measurement. Here, homogenizations were performed individually into 50 µL buffer for workers, into 200µL for queens, with micropilons. To remove tissues debris, the homogenized samples were centrifuges at 4°C. They were then diluted by 1:200 before reading plate, to stay in the standard range. Results were normalized using sample protein content with Bradford method (BioRad).
