## Supplementary Figure S1 for "Mitochondrial maintenance is involved in the exceptional longevity of reproductive queens of the eusocial ant *Lasius niger*"

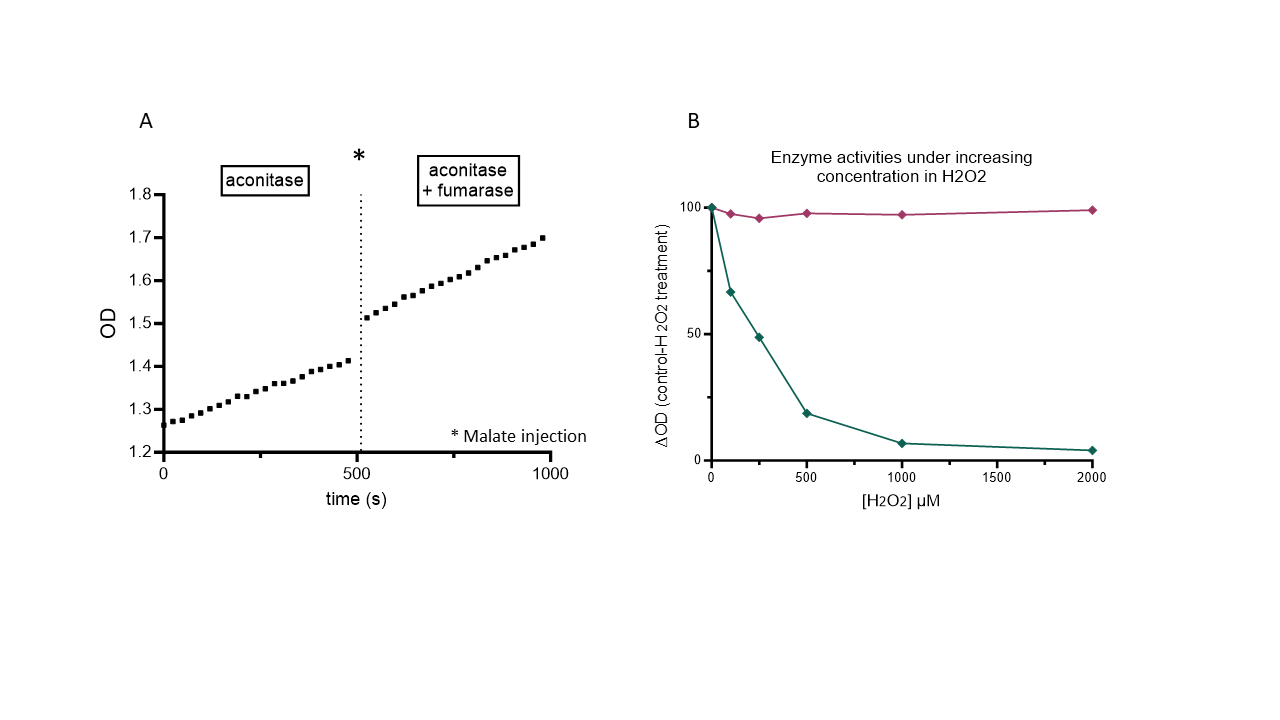

Figure S1: Procedure for measuring in a well. The first kinetic is initiated by adding isocitrate (t0), and the second by adding malate (A). Aconitase and fumarase activities in the same homogenate after increasing addition of H_2_O_2_, in order to verify aconitase sensitivity to oxidative stress in the model. Aconitase activity is in green, fumarase activity in purple (B).
