## Supplementary figures and images for "Mitochondrial maintenance is involved in the exceptional longevity of reproductive queens of the eusocial ant *Lasius niger*"

### Supplementary Figure S2

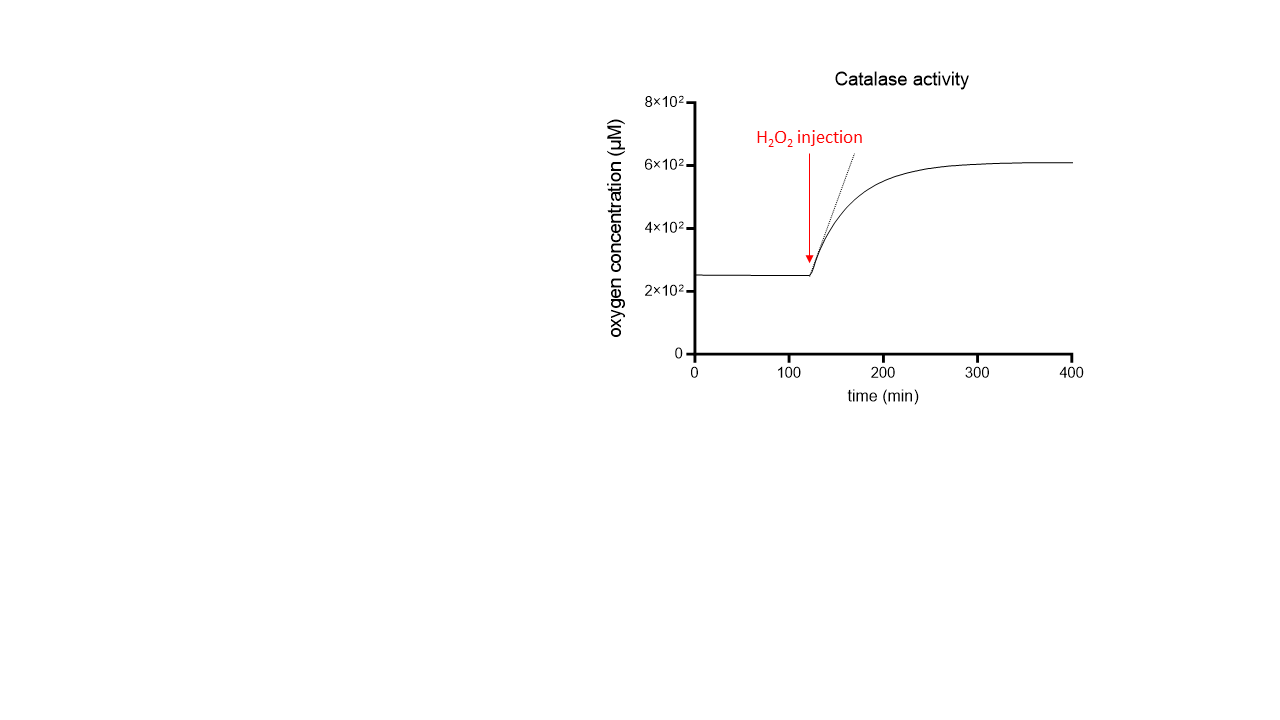

Figure S2: Catalase activity was measured with a Clark electrode by H_2_0_2_ to O_2_ conversion
