## Supplementary Figure S3 for "Mitochondrial maintenance is involved in the exceptional longevity of reproductive queens of the eusocial ant *Lasius niger*"

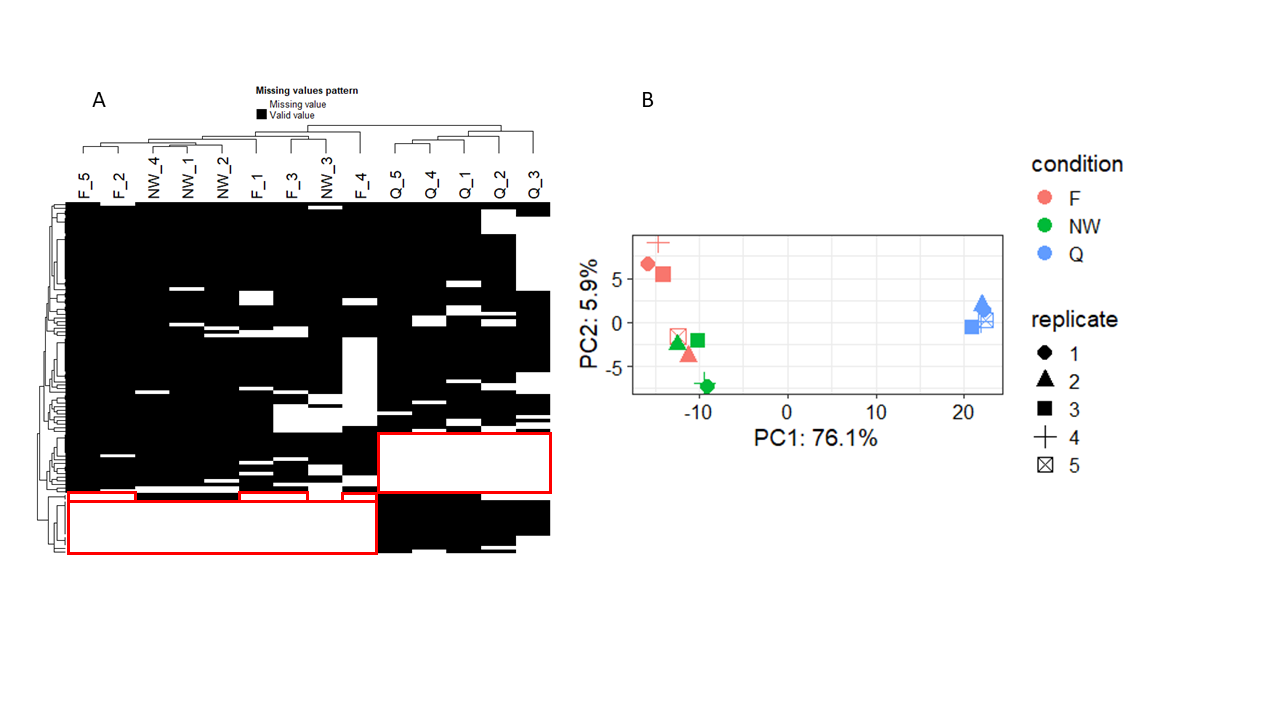

Figure S3: Heatmap of proteins (black) with missing values (white). Missing values not at random are framed in red (A). PCA analysis on previous published data by Quque et al [3] retaining only mitochondrial proteins: individuals plot (B). Q: queen, NW: nest-worker, F: forager
