## Supplementary Figure S4 for "Mitochondrial maintenance is involved in the exceptional longevity of reproductive queens of the eusocial ant *Lasius niger*"

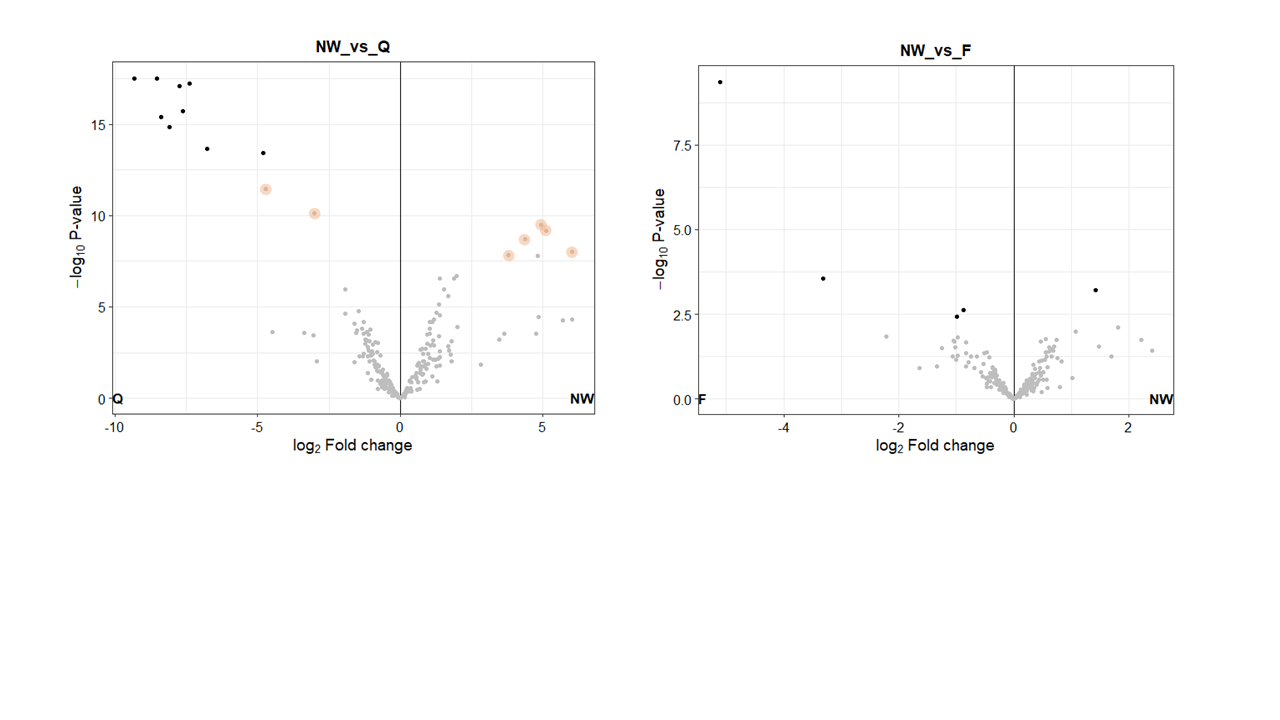

Figure S4: (A) Volcano plots obtained with queen caste as control. Significant protein fold changes compared to the nest-worker group are in bold. Proteins highlighted in orange were significantly over- or under-expressed in the forager group. (B) Volcano plot between the worker groups with foragers as control.
