## Supplementary Figure S5 for "Mitochondrial maintenance is involved in the exceptional longevity of reproductive queens of the eusocial ant *Lasius niger*"

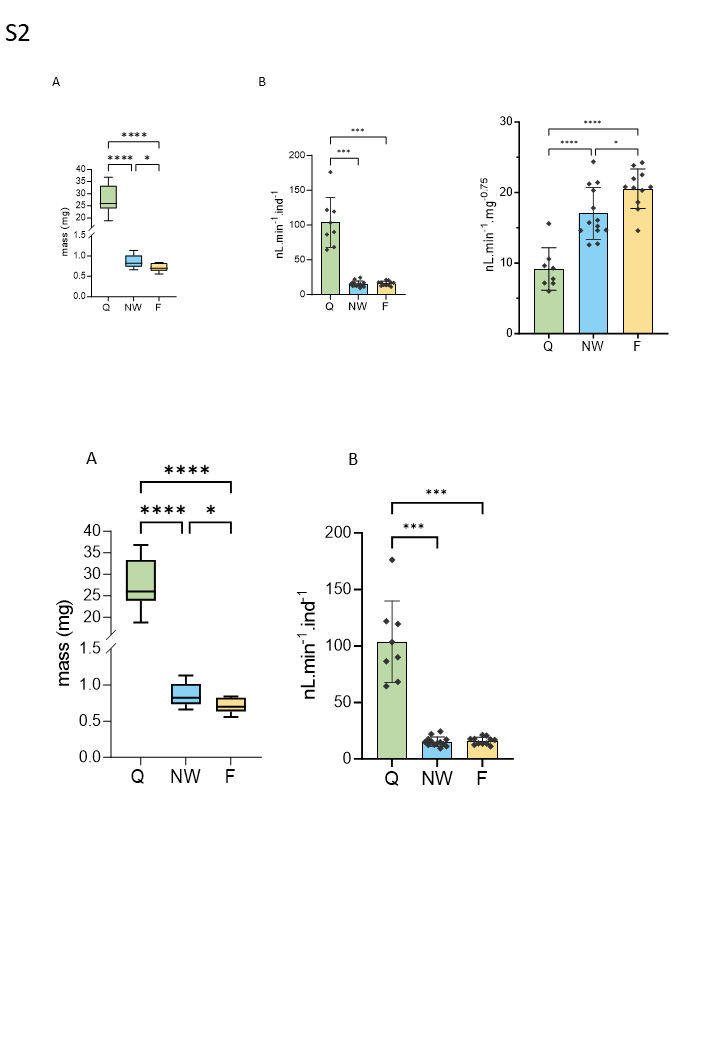

Figure S5: Ant mass measured for single queen or worker pools (A), and oxygen consumption reported per individual. Q: queen, NW: nest-worker, F: forager. Values are mean ± SD. ****P <0.0001, P***<0.0005, P*<0.05
