## Supplementary Figure S6 for "Mitochondrial maintenance is involved in the exceptional longevity of reproductive queens of the eusocial ant *Lasius niger*"

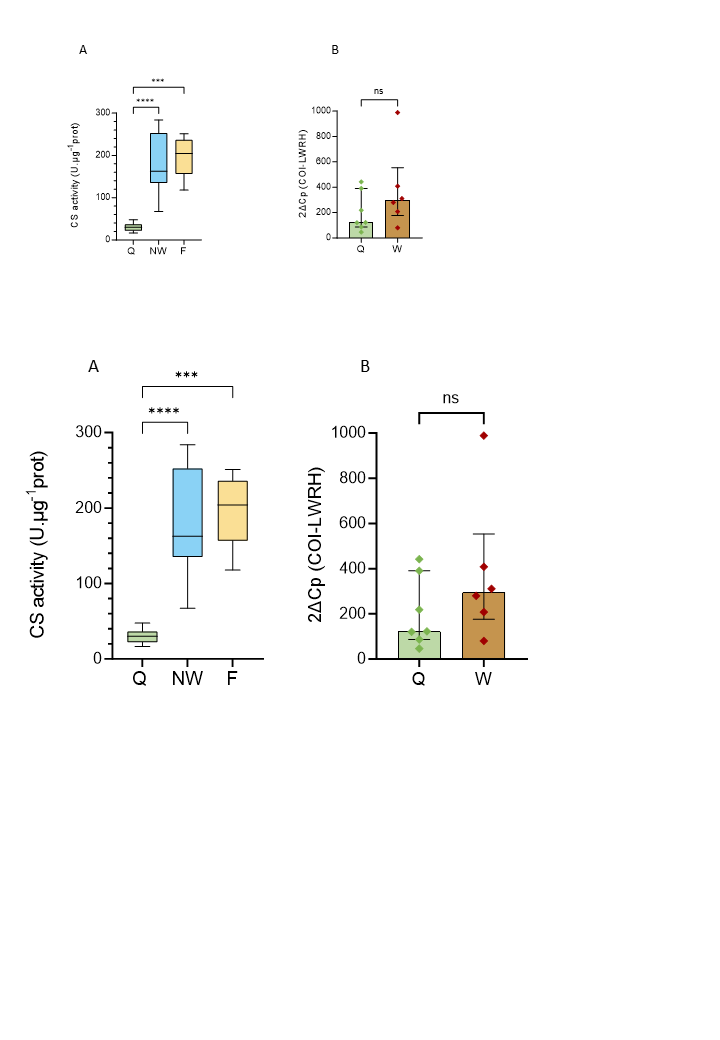

Figure S6: Mitochondrial densities measured by citrate synthase activity (A) relative mtDNA copy number of COI by qPCR (B), on whole body. Values are mean ± SD in A, median (IQR) in B. Q: queen, NW: nest-worker, F: forager. ****P <0.0001, P***<0.001.
