## Supplementary Figure S7 for "Mitochondrial maintenance is involved in the exceptional longevity of reproductive queens of the eusocial ant *Lasius niger*"

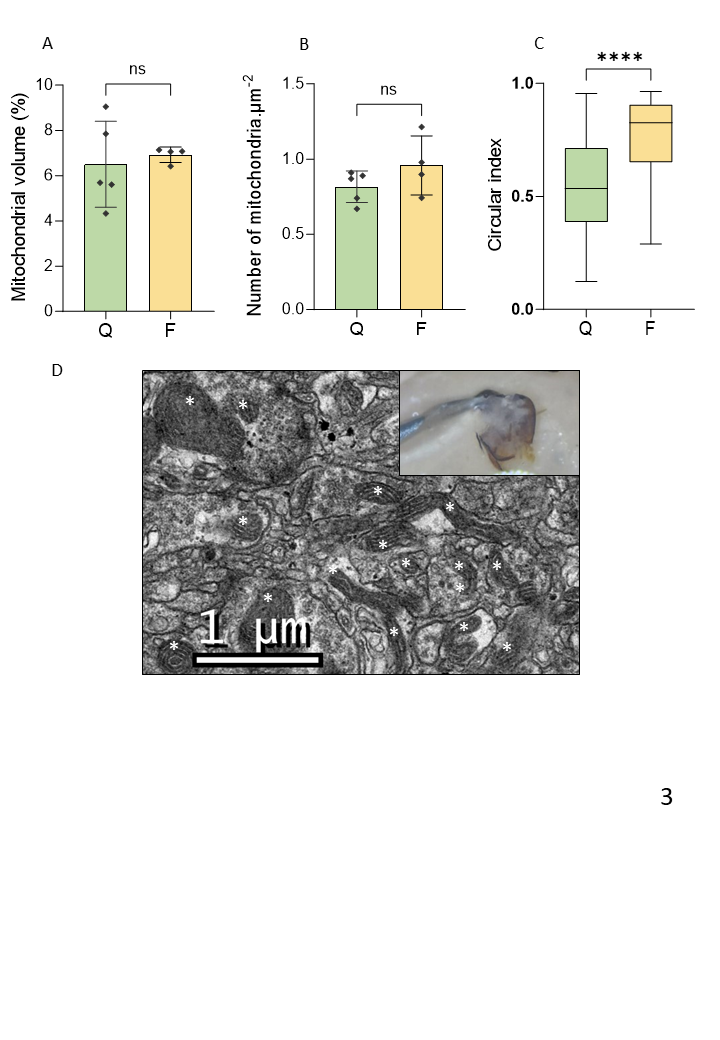

Figure S7: Mitochondrial volume measured by electron microscopy in the brain tissues (A), numbers of mitochondria per section were reported in (B), and finally circular index in a boxplot for a relative comparison of fission/fusion events (C). Values are mean ± SD for A and B, the boxplot C represents the median, quartiles and the range of values. Q: queen, F: forager. ****P <0.0001. For analyses, ants were dissected in fixative solution to avoid at maximum mitochondrial oxidation. Here the forager brain dissection, whose mitochondria are indicated with the white asterisks under microscopy (D).
