## Supplementary Figure S8 for "Mitochondrial maintenance is involved in the exceptional longevity of reproductive queens of the eusocial ant *Lasius niger*"

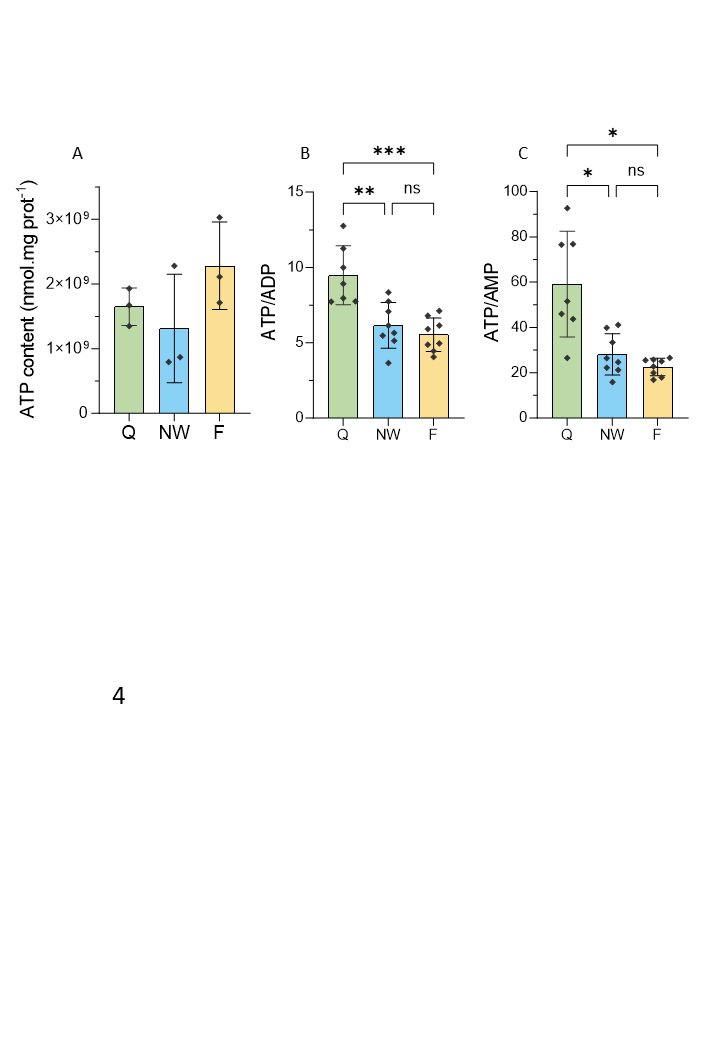

Figure S8: ATP content obtained via bioluminescence (A), ATP/ADP ratio (B) and ATP/AMP ratio (C) obtained from the AEC. Values are mean ± SD for A and B, median (IQR) in C. Q: queen, NW: nest-worker, F: forager. P***<0.0005, P**<0.005, P*<0.05.
