## Supplementary Figure S9 for "Mitochondrial maintenance is involved in the exceptional longevity of reproductive queens of the eusocial ant *Lasius niger*"

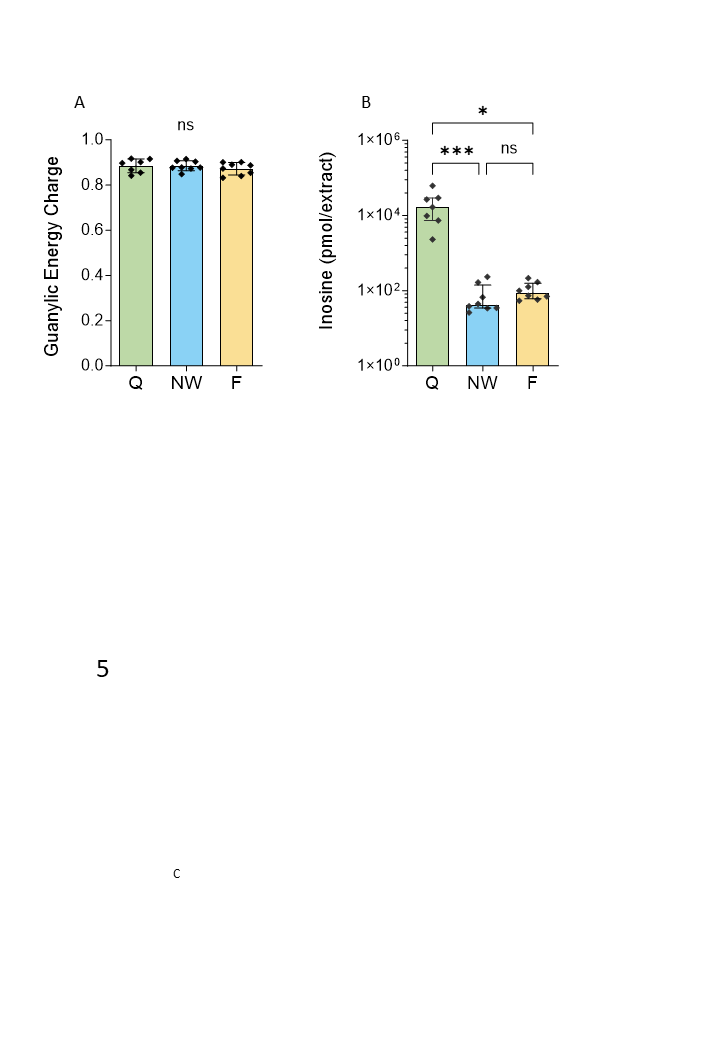

Figure S9: Guanylic Energy Charge (A), and inosine concentration on homogenates (B). Values are mean ± SD for A, median (IQR) in B. Q: queen, NW: nest-worker, F: forager. P***<0.0005, P*<0.05
