## Supplementary Figure S10 for "Mitochondrial maintenance is involved in the exceptional longevity of reproductive queens of the eusocial ant *Lasius niger*"

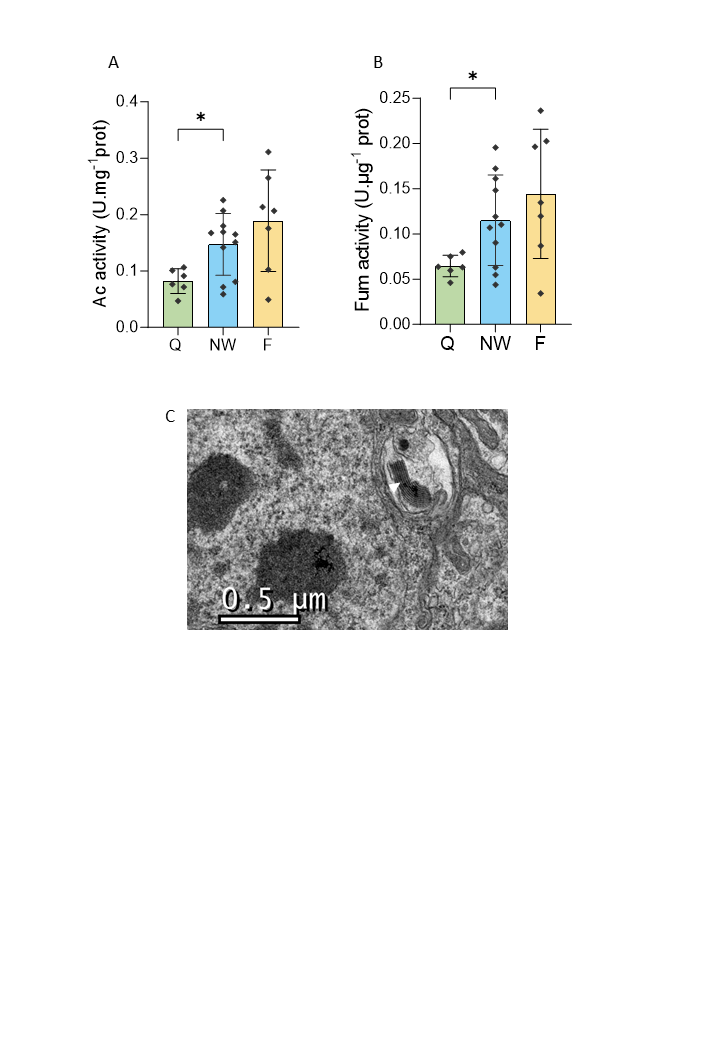

Figure S10: Aconitase (A) and fumarase activities (B) measured in a same well. Electron microscopy allowed to observe lipofuscin accumulation in ant’s brain (C), indicated with the white arrow. Values are median (IQR). Q: queen, NW: nest-worker, F: forager. P*<0.05.
