## Supplementary Figure S11 for "Mitochondrial maintenance is involved in the exceptional longevity of reproductive queens of the eusocial ant *Lasius niger*"

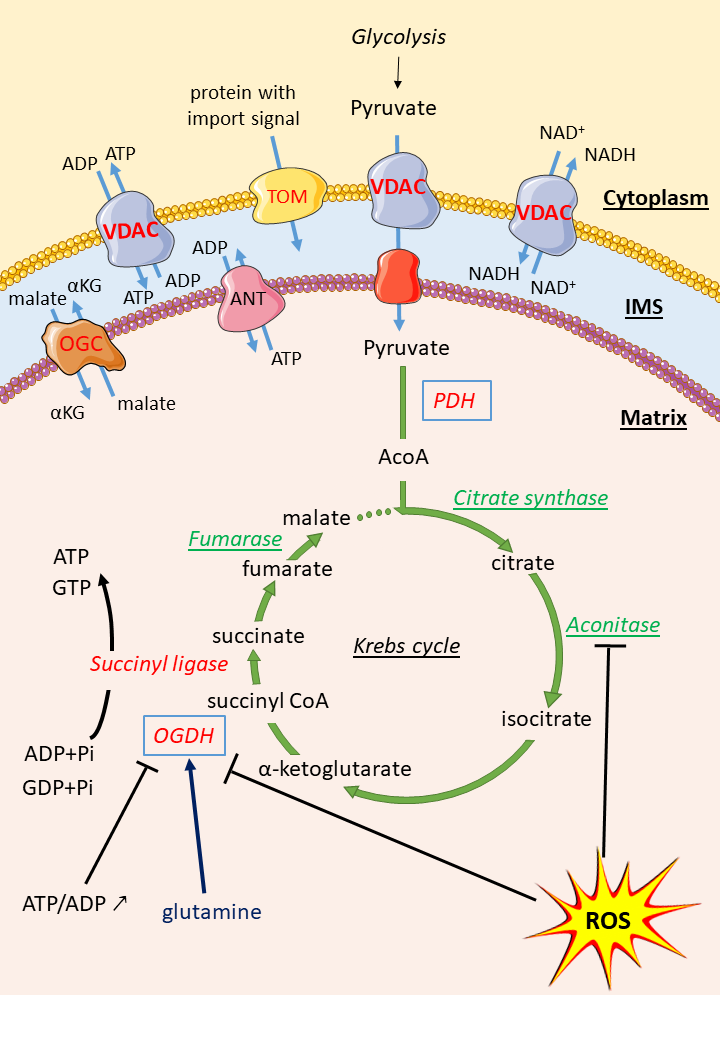

Figure S11: mitochondrial membrane flux and flux in the Krebs cycle. Proteins with at least one subunit showing a significant fold change in favour of queens in proteomics are highlighted in red. The Krebs cycle enzymes underlined in green were assessed in this paper and exhibit lower activity in queens compared to at least one group of workers. VDAC is a membrane protein that facilitates the exchange of metabolites of various types between cytoplasm and mitochondrial intermembrane space, contributing to the regulation of the Krebs cycle and energy production. The ANT translocase allows the entry of ADP into the mitochondrial matrix in exchange for ATP. PDH and OGDH are key **players** controlling the large majority of carbon flow into the Krebs cycle, derived from pyruvate and glutamine, respectively. KGDH is particularly sensitive to ROS (especially H_2_O_2_), as is aconitase, whose inactivation serves as a marker of oxidative stress when the aconitase/fumarase ratio decreases. PDH: pyruvate dehydrogenase, OGDH: 2-oxoglutarate dehydrogenase E1, VDAC: voltage-dependent anion channel proteins, OMT: oxoglutarate-malate translocator, TOM: translocase of the outer membrane, AcoA: acetyl coenzyme A, IMS: intermembrane space, αKG: alpha-ketoglutarate, ROS: reactive oxygen species (H_2_O_2_).
