## Supplementary Table S1 for "Mitochondrial maintenance is involved in the exceptional longevity of reproductive queens of the eusocial ant *Lasius niger*"

Table S1: Number of ants (or pools of ants) used for each analysis. The counts in the red row (enzymatic activities) correspond to the same ants as those in the row above, but with a reduced number of specimens.

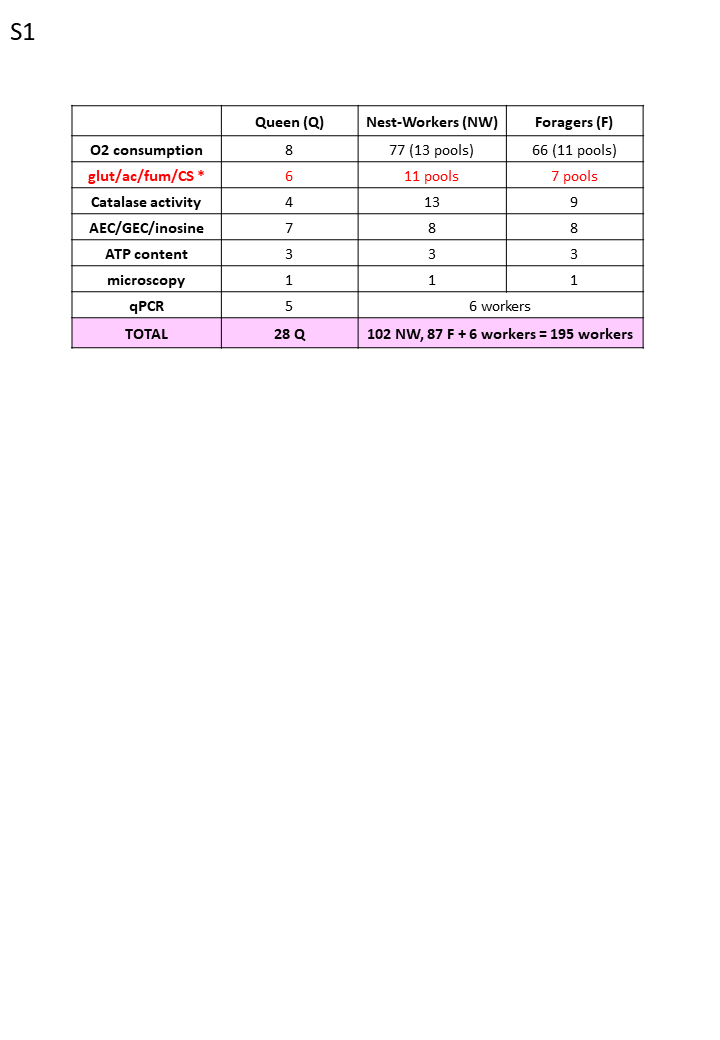
