## Supplementary Table S2 for "Mitochondrial maintenance is involved in the exceptional longevity of reproductive queens of the eusocial ant *Lasius niger*"

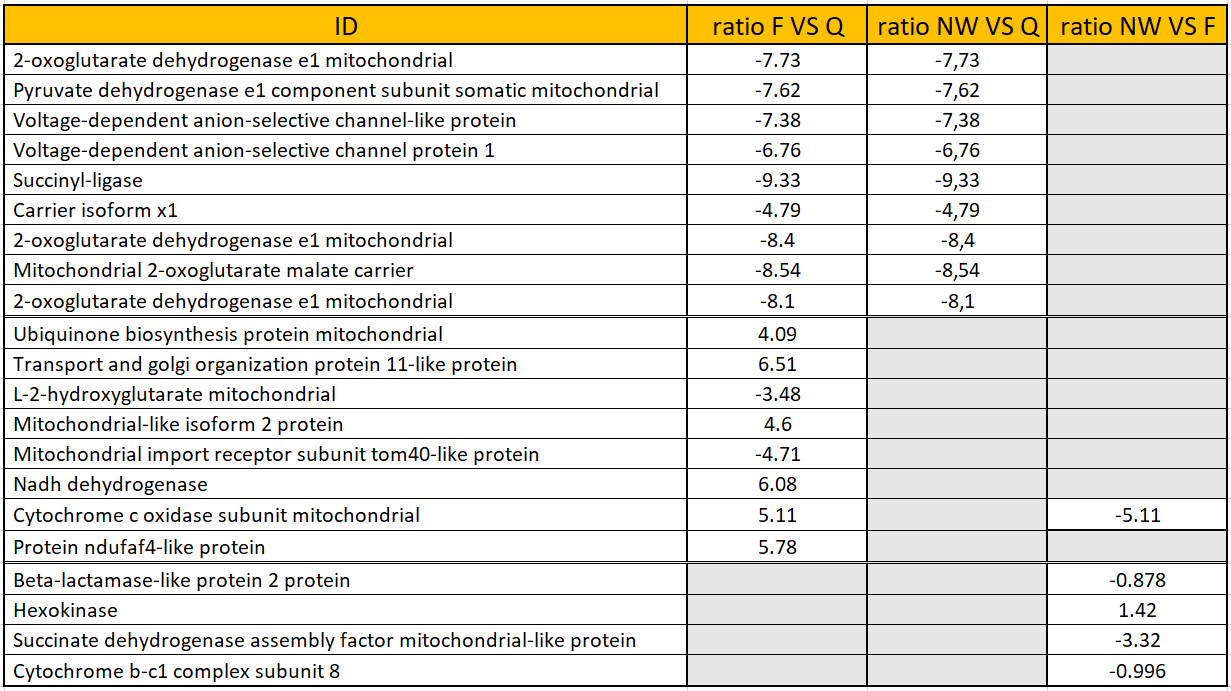
Table S2: Significant results from the proteomic analyses (P-value < 0.05). In the first analysis, the queens were used as the reference. Subsequently, foragers were used as the reference for comparisons among workers. A negative ratio indicates that proteins are more abundant in the reference group.
